# ATP synthase-associated CHCH domain proteins are critical for mitochondrial function in *Toxoplasma gondii*

**DOI:** 10.1101/2023.02.02.526833

**Authors:** Madelaine M. Usey, Diego Huet

## Abstract

Coiled-coil-helix-coiled-coil-helix (CHCH) domains consist of two pairs of cysteine residues that are oxidized to form disulfide bonds upon mitochondrial import. Proteins containing these domains play important roles in mitochondrial ultrastructure and in the biogenesis, function, and stability of electron transport chain complexes. Interestingly, recent investigations of the *Toxoplasma gondii* ATP synthase identified subunits containing CHCH domains. As CHCH domain proteins have never been found in any other ATP synthase, their role in *T. gondii* was unclear. Using conditional gene knockdown systems, we show that two *T. gondii* ATP synthase subunits containing CHCH domains are essential for the lytic cycle as well as stability and function of the ATP synthase. Further, we illustrated that knockdown disrupts multiple aspects of mitochondrial morphology. Mutation of key residues in the CHCH domains also caused mislocalization of the proteins. This work provides insight into the divergent aspects of the apicomplexan ATP synthase, which could uncover future drug targets.

## INTRODUCTION

The adenosine triphosphate synthase (ATP synthase) is a membrane-bound multisubunit enzyme responsible for generating cellular energy in the form of ATP. This complex consists of two distinct parts: the membrane-bound F_0_ portion and the knob-like catalytic F_1_ portion, which extends away from the membrane. The F_0_ and F_1_ portions are functionally linked via a central and peripheral stalk (Rubinstein, Walker, and Henderson 2003). Within eukaryotic mitochondria, the complexes of the electron transport chain (ETC) generate a proton gradient across the inner mitochondrial membrane. The gradient can be converted into rotation of the ATP synthase central stalk by proton translocation through the F_0_ portion of the enzyme. This rotation, which is counteracted by the peripheral stalk, causes conformational changes that allow for the generation of ATP from ADP and inorganic phosphate via the process of oxidative phosphorylation: a well-characterized and conserved process (Nakamoto, Baylis Scanlon, and Al-Shawi 2008; Sobti et al. 2021; Weber 2007). While the general architecture and subunit composition of the ATP synthase is similarly conserved among a wide array of organisms including plants, bacteria, and mammals (John E. Walker 2013), relatively little is known about the ATP synthase in members of the apicomplexan phylum.

Apicomplexan parasites are a large phylum of eukaryotic pathogens that impose a significant burden on global public health. Previous work in *T. gondii* and *Plasmodium spp*., the causative agents of toxoplasmosis and malaria, respectively, has illustrated that these parasites can shift their reliance on glycolysis or oxidative phosphorylation for ATP production to fit varying metabolic needs faced throughout their complex life cycles (J. I. MacRae et al. 2012; James I. MacRae et al. 2013). As such, studies in the mouse model of malaria, *Plasmodium berghei*, found that while disruption of the ATP synthase was only modestly detrimental to asexual blood stages, it completely blocked sexual replication during the insect stages of the life cycle (Sturm et al. 2015). While *T. gondii* tachyzoites also exhibit metabolic flexibility (Blume et al. 2009; Fleige et al. 2008; Shukla et al. 2018), an intact ATP synthase was shown to be essential for tachyzoite survival, and it has been estimated that more than 80% of ATP in egressed tachyzoites is generated via the TCA cycle under normal conditions (J. I. MacRae et al. 2012; Huet et al. 2018).

In addition to their role in energy production, another important aspect of mitochondrial ATP synthases is their ability to form oligomeric complexes. In yeast and mammals, ATP synthase dimers have been shown to be critical for cristae formation by inducing curvature of the inner mitochondrial membrane, a process that is important for maintaining mitochondrial membrane potential and promoting efficient ATP production (Blum et al. 2019; Strauss et al. 2008; Bornhövd et al. 2006). The *T. gondii* ATP synthase was found to assemble into dimers via an extensive interface mediated primarily by apicomplexan-specific subunits (Muhleip et al. 2021). With a molecular weight of ∼1860 kDa (Maclean et al. 2021; Muhleip et al. 2021), the *T. gondii* ATP synthase dimer is significantly larger than the ∼1200 kDa dimer of yeast and mammals (Lau, Baker, and Rubinstein 2008; J. E. Walker et al. 1991), a trend which has been observed with most *T. gondii* respiratory complexes (Usey and Huet 2022).

Beyond its significantly larger size, multiple studies have also found that the subunit composition of the *T. gondii* ATP synthase is highly divergent compared to other model organisms (Huet et al. 2018; Salunke et al. 2018; Muhleip et al. 2021). In all, the *T. gondii* ATP synthase consists of 32 subunits, of which only 15 are canonical and have homologs in other phyla (Muhleip et al. 2021). The other 17 subunits are generally conserved only among mitochondriate apicomplexans. However, some are also found within other organisms of the Myzozoan clade, a phylogenetic group which includes chromerids, perkinsozoa, and apicomplexans. Further, several canonical subunits have extended apicomplexan-specific domains with functions that remain to be determined (Muhleip et al. 2021).

Though one of the divergent *T. gondii* ATP synthase subunits has been studied (Muhleip et al. 2021), the rest lack functional characterization. Interestingly, three of the novel phylum-specific F_0_ subunits have been found to contain putative coiled-coil-helix-coiled-coil-helix (CHCH) domains (Huet et al. 2018; Muhleip et al. 2021). CHCH domains consist of sets of cysteine residues as CX_9_C motifs, in which the X represents any other amino acid, within separate α-helices of the protein (Modjtahedi et al. 2016). In other organisms, proteins containing CHCH domains are often found as subunits of multi-protein complexes within the mitochondrion. Many proteins containing these domains are subunits of complexes I, III, and IV of the ETC where they are important for structure, assembly, optimal function, and chaperoning copper ions (Modjtahedi et al. 2016; Cavallaro 2010; S. Longen et al. 2009). CHCH-domain proteins are also found to play roles in mitochondrial lipid transport (Potting et al. 2010), as part of the mitochondrial cristae organizing system (MICOS) (Manjula Darshi et al. 2011), and as part of the mitochondrial ribosome (Sebastian Longen et al. 2014).

The three proteins containing CHCH domains that were identified as subunits of the *T. gondii* ATP synthase also appear to be conserved in other apicomplexans (Huet et al. 2018; Muhleip et al. 2021). However, because CHCH domain proteins have never before been observed as part of the ATP synthase in other organisms, it is unclear what role they play in apicomplexans. In this study, we characterized two of the *T. gondii* ATP synthase subunits containing CHCH domains: TGGT1_258060 (ATPTG8) and TGGT1_285510 (ATPTG9). Using conditional gene knockdown systems, we show that both genes are essential for the *T. gondii* lytic cycle and are critical for ATP synthase structural stability, oxidative phosphorylation, mitochondrial membrane potential maintenance, proper mitochondrial morphology, and cristae density. Further, we demonstrated that the CHCH domain cysteine residues in both ATPTG8 and ATPTG9 are essential for mitochondrial localization of both proteins. This study deepens our understanding of the apicomplexan ATP synthase and of the role played by the divergent subunits conserved throughout these pathogens.

## RESULTS

### *T. gondii* ATP synthase subunits containing CHCH domains are essential for the lytic cycle

ATPTG8 and ATPTG9 were originally found to be associated with the *T. gondii* ATP synthase via immunoprecipitation and cryo-electron microscopy studies (Huet et al. 2018; Salunke et al. 2018; Muhleip et al. 2021). ATPTG8 contains a single CHCH domain towards the C terminus of the gene. In contrast, ATPTG9 contains two CHCH domains (Figure 1A), which occurs in only about 8% of CHCH-domain proteins (Cavallaro 2010).

**Figure 1:**
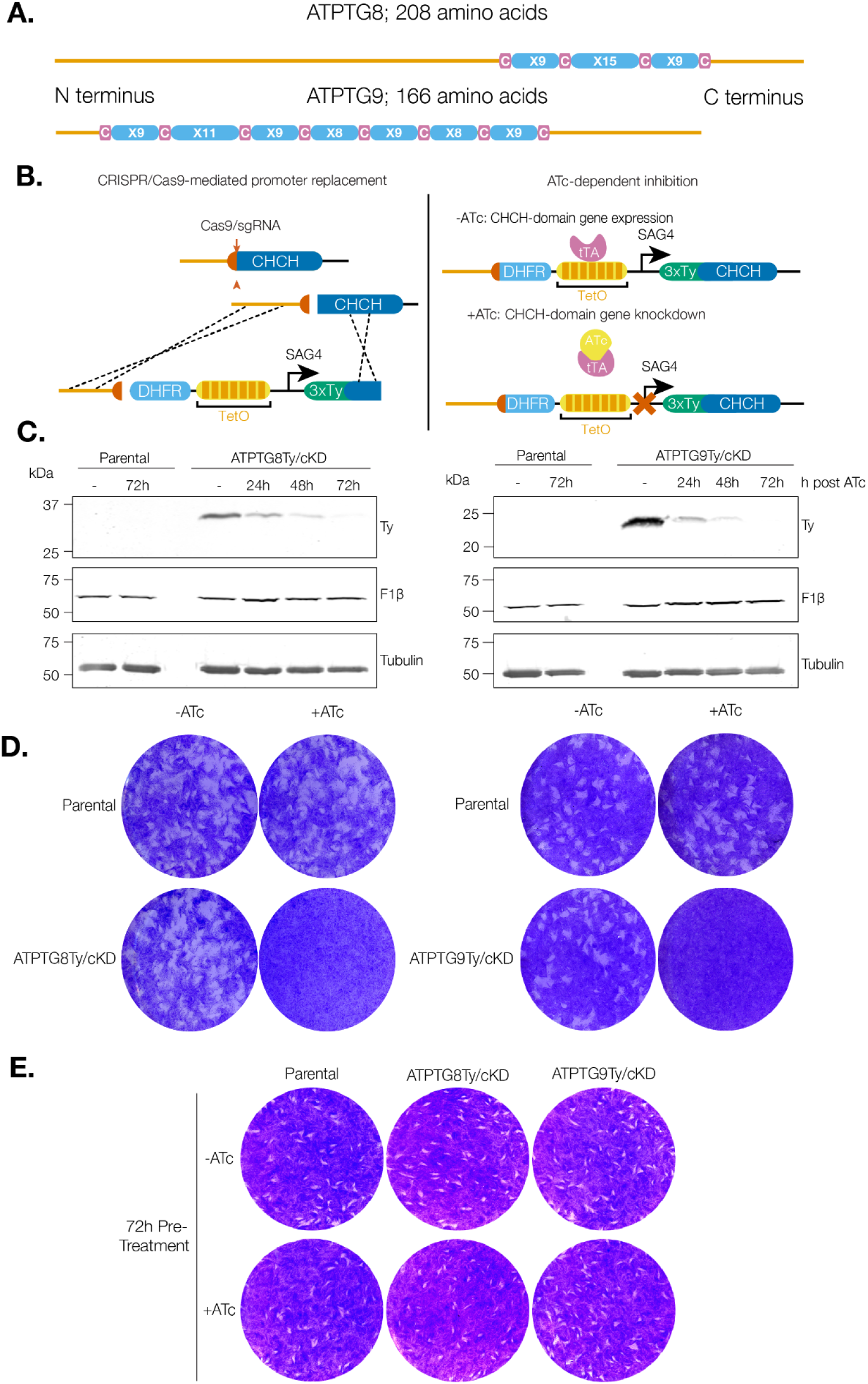
CHCH domain proteins associated with the *T. gondii* ATP synthase are essential. **A**. Schematic representation of the CHCH domain size and location in ATPTG8 and ATPTG9. C represents cysteine residues and X represents any other amino acid residue. **B**. Schematic representation of the strategy to generate ATPTG8 and ATPTG9 conditional knockdown strains (ATPTG8Ty/cKD and ATPTG9Ty/cKD). A pyrimethamine-resistant dihydrofolate reductase (DHFR) cassette, the *T. gondii* SAG4 promoter, a tetracycline-inducible operator, and an in-frame 3xTy epitope tag were inserted into the promoter region of either ATPTG8 or ATPTG9 via CRISPR/Cas9 and homology-directed repair. When anhydrotetracycline (ATc) is added to the culture medium, the tetracycline transactivator (tTA) expressed in the TATi/Δku80 parental line is no longer able to bind TetO, resulting in gene knockdown. **C**. Lysates from parental, ATPTG8Ty/cKD, and ATPTG9Ty/cKD parasites were prepared following treatment with ATc or vehicle control for the indicated amounts of time. Samples were separated via SDS-PAGE then probed with antibodies against Ty, F1β, or tubulin. **D**. Plaque assay of parental, ATPTG8Ty/cKD, or ATPTG9Ty/cKD parasites grown undisturbed on an HFF monolayer in the presence of ATc or vehicle control for 7-8 days. **E**. Plaque assay of parental, ATPTG8Ty/cKD, or ATPTG9Ty/cKD parasites pre-treated with ATc or vehicle control for 72h then grown undisturbed on an HFF monolayer for 7-8 days.

To begin characterizing the roles of ATPTG8 and ATPTG9, we utilized a conditional gene knockdown system, as both genes were previously predicted to be essential for the *T. gondii* lytic cycle (Sidik et al. 2016). Using CRISPR/Cas9 and homology-mediated repair, we replaced the promoter of each gene in the TATi/Δku80 strain with the SAG4 promoter, a tetracycline-inducible operator (TetO), and an in-frame N-terminal 3xTy epitope tag (Figure 1B) (Sheiner et al. 2011; Meissner et al. 2001), thus creating ATPTG8Ty/cKD and ATPTG9Ty/cKD parasite lines. With this conditional gene knockdown system, when anhydrotetracycline (ATc) is added to the culture medium, the tetracycline transactivator (tTA) expressed in the TATi/Δku80 parental line is no longer able to bind TetO, inhibiting gene transcription. Once clonal populations for each line were obtained, each strain was treated with vehicle control (ethanol) or ATc for 24, 48, or 72 hours. Detection of ATPTG8Ty and ATPTG9Ty by Western blot shows expression of Ty-tagged proteins at the correct molecular weight in the absence of ATc and depletion of Ty signal over 72 hours of ATc treatment (Figure 1C).

To confirm the essentiality of ATPTG8 and ATPTG9 for the *T. gondii* lytic cycle, we allowed ATPTG8Ty/cKD, ATPTG9Ty/cKD, and parental parasites to invade and replicate in a human foreskin fibroblast (HFF) monolayer undisturbed for 7-8 days in the presence of ATc or vehicle control. While the ability of the parental line to form plaques was unaffected by the addition of ATc, neither ATPTG8Ty/cKD nor ATPTG9Ty/cKD parasites were able to form any plaques in the presence of ATc (Figure 1D), affirming that they are essential for the lytic cycle. However, when ATPTG8Ty/cKD and ATPTG9Ty/cKD parasites were pre-treated with ATc for 72 hours before addition to the HFF monolayer with normal media, they were able to recover and form plaques (Figure 1E). This experiment establishes that the phenotypic abnormalities discussed below are not the result of parasite death at this time point, but instead reflect specific defects resulting from the loss of ATPTG8 or ATPTG9 subunits.

### ATP synthase subunits containing CHCH domains are required for structural stability and function of the complex

As ATPTG8 and ATPTG9 have previously been determined to be subunits of the *T. gondii* ATP synthase (Huet et al. 2018; Muhleip et al. 2021; Maclean et al. 2021), we next wanted to determine their role in both the structure and function of this key enzymatic complex. To first investigate their structural roles, we utilized blue native-PAGE (BN-PAGE). Using an antibody against the β subunit of the ATP synthase, we were able to observe a strong band above 1700 kDa in the parental strain grown in the presence of ATc or vehicle control, as well in the conditional knockdown strains grown in vehicle control for 72h (Figure 2A). This molecular weight corresponds to the reported ∼1860 kDa mass of the *T. gondii* ATP synthase dimer (Muhleip et al. 2021; Maclean et al. 2021). A band of intermediate weight (∼123-215 kDa) could also be observed in these samples, likely indicating either electrophoretic degradation or an ATP synthase assembly intermediate. Similarly, the lowest molecular weight band observed (below 123 kDa), likely represents single β subunits. When ATPTG8Ty/cKD and ATPTG9Ty/cKD parasites were treated with ATc for increasing amounts of time, the dimeric form of the ATP synthase was eventually lost, leading to increased ATP synthase intermediate forms and free β subunit. Interestingly, slightly more intermediates and free β subunit were consistently observed in ATPTG8Ty/cKD parasites not treated with ATc, suggesting that either modified expression of the ATPTG8 gene due to promoter replacement or the 3xTy tag at the N terminus of the protein could have caused a slight change in ATP synthase dimerization (Figure 2A). Overall, these experiments illustrate that ATPTG8 and ATPTG9 are critical for ATP synthase structural stability in *T. gondii*.

**Figure 2:**
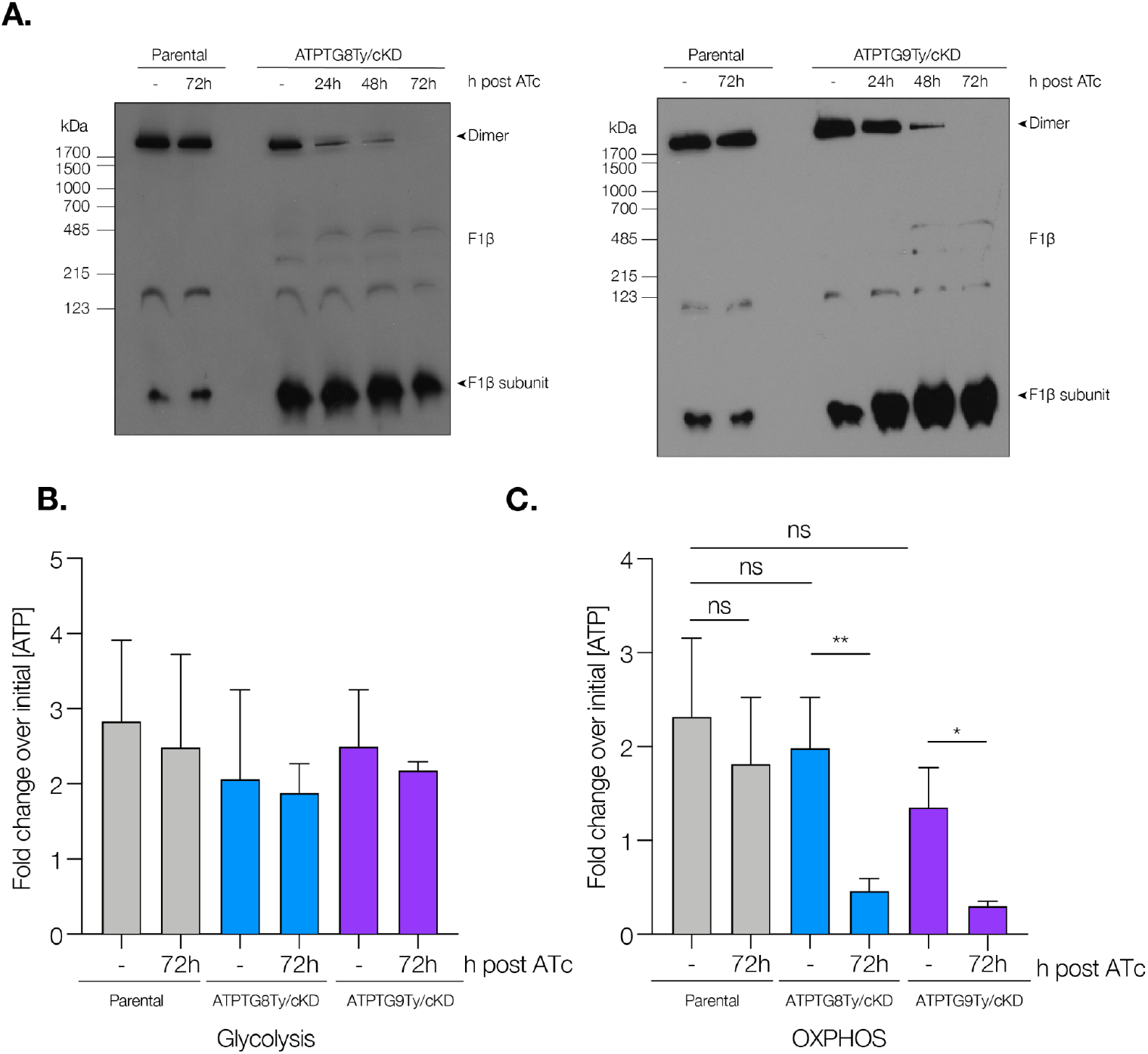
ATP synthase subunits containing CHCH-domains are required for ATP synthase stability and function. **A**. Lysates from parental, ATPTG8Ty/cKD, and ATPTG9Ty/cKD parasites were prepared following treatment with ATc or vehicle control (-) for the indicated time points, then resolved by blue native PAGE (BN-PAGE) and probed for F1β. (**B**,**C)** Relative ATP concentrations of parental, ATPTG8Ty/cKD, and ATPTG9Ty/cKD parasites were measured following a 72h treatment with ATc or vehicle control. Following mechanical release, parasites were incubated for 1 hour with 5mM 2-deoxy-D-glucose (2-DG) combined with **(B)** 25mM glucose to promote glycolysis or **(C)** 2mM glutamine to promote oxidative phosphorylation. To determine relative concentrations, ATP levels for each condition were normalized to the initial ATP readout of each strain. Data represent mean ± SD of 6 independent replicates for parental strains, and 3 independent replicates for ATPTG8Ty/cKD and ATPTG9Ty/cKD strains. For one independent replicate of ATPTG8Ty/cKD, only two technical replicates were used for glucose and glutamine treatment; ns = not significant, p = 0.001 to 0.01: **, p = 0.01 to 0.05: *, unpaired, two-tailed t-test).

Since ATP synthase structure was severely disrupted following CHCH domain gene knockdown, we wanted to see whether this was associated with a concurrent disruption in ATP synthase activity. To study the metabolic effects of CHCH domain gene knockdown, we utilized a previously published metabolic assay (Huet et al. 2018). Briefly, intracellular ATPTG8Ty/cKD, ATPTG9Ty/cKD, and parental parasites were treated with ATc or vehicle for 72h before addition of the glycolysis inhibitor 2-deoxy-D-glucose (2-DG). Following this treatment, sufficient glucose to promote glycolysis or sufficient glutamine to promote oxidative phosphorylation was added to the parasites. ATP levels were measured and relative ATP concentrations were determined for each treatment condition through comparison to initial ATP levels. Though ATP produced via glycolysis was not significantly different between strains regardless of ATc treatment (Figure 2B), there were significant decreases in the ATP produced via oxidative phosphorylation when ATPTG8Ty/cKD and ATPTG9Ty/cKD parasites were treated with ATc for 72h (Figure 2C). In summary, knockdown of these ATP synthase subunits containing CHCH domains decreases ATP production via the ATP synthase, but not by glycolysis.

As ATP synthase function is closely tied to mitochondrial membrane potential, we also utilized MitoTracker staining and flow cytometry to investigate how ATPTG8 or ATPTG9 knockdown affects mitochondrial membrane potential (Figure S1). With this method, we observed that ATPTG9 knockdown over 72 hours of ATc treatment resulted in a significant membrane potential depolarization compared to the parental strain. Though ATPTG8 knockdown showed decreased membrane potential compared to parental parasites, this difference was not statistically significant (p=0.068) (Figure S1). Therefore, these results suggest that ATPTG8 and ATPTG9 are important for proper maintenance of the mitochondrial membrane potential.

### CHCH domain proteins are important for *T. gondii* mitochondrial morphology and volume

In *T. gondii*, as well as in humans, defects in ATP synthase assembly can lead to distorted, swollen, and abnormally shaped mitochondria (Galber et al. 2021; Huet et al. 2018; Dautant et al. 2018). To investigate whether similar defects are observed upon knockdown of ATPTG8 and ATPTG9, we utilized immunofluorescence microscopy. Mitochondrial morphology was observed via staining against the outer mitochondrial membrane protein TOM40 (van Dooren et al. 2016). In intracellular ATPTG8Ty/cKD and ATPTG9Ty/cKD parasites treated with vehicle control, as well as in parental parasites regardless of ATc treatment, TOM40 staining consistently showed a lasso-shaped signal (Figure 3A). This is in line with previous observations illustrating that most intracellular *T. gondii* parasites exhibit lasso-shaped mitochondria (Ovciarikova et al. 2017). However, as ATPTG8Ty/cKD and ATPTG9Ty/cKD parasites are treated with ATc for increasing amounts of time, mitochondria appear increasingly fragmented and abnormally shaped as indicated by the TOM40 staining (Figure 3A). These data indicate that loss of ATPTG8 and ATPTG9 critically affects mitochondrial health and general morphology.

**Figure 3:**
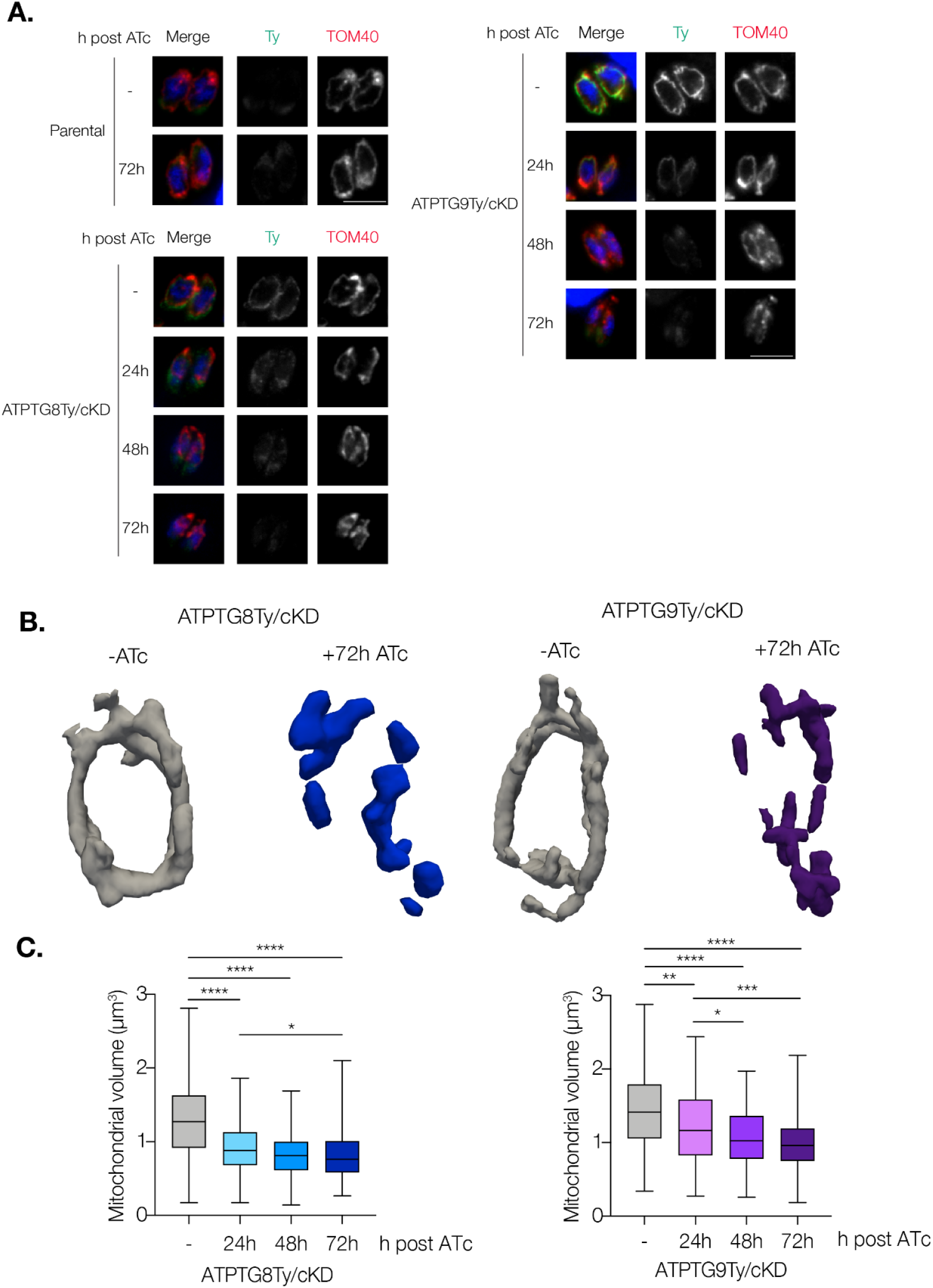
CHCH domain proteins are critical for mitochondrial morphology and volume. **A**. Intracellular parental, ATPTG8Ty/cKD, or ATPTG9Ty/cKD parasites were fixed and stained for Ty (green) and the mitochondrial marker TOM40 (red) following addition of ATc or vehicle control (-) for the indicated time points. Representative images from 3 independent replicates. Scale bar: 5µm. **(B**,**C)** Mitochondria from intracellular ATPTG8Ty/cKD and ATPTG9Ty/cKD parasites expressing SOD2-GFP and IMC1-TdTomato were analyzed via MitoGraph. **B**. Representative 3D MitoGraph projections of mitochondria from ATPTG8Ty/cKD and ATPTG9Ty/cKD parasites following treatment with ATc or vehicle control (-) for 72h. **C**. Quantification of mitochondrial volumes from ATPTG8Ty/cKD and ATPTG9Ty/cKD parasites after treatment with ATc or vehicle control (-) for the indicated times. At least 100 vacuoles were measured for each condition over 3 independent replicates for ATPTG8Ty/cKD or 2 independent replicates for ATPTG9Ty/cKD. Unpaired, two-tailed t-test (p < 0.0001: ****, p = 0.0001 to 0.001: ***, p = 0.001 to 0.01: **, p = 0.01 to 0.05: *).

Though immunofluorescence can be useful for investigating gross mitochondrial defects, we wanted to delve deeper into the mitochondrial abnormalities we observed upon ATP synthase-associated CHCH domain protein knockdown. To do so, we utilized mitochondrial volume analysis, a technique that has previously been used in both yeast and *T. gondii* to monitor mitochondrial volume in three dimensions and quantify changes in mitochondrial volume (Viana, Lim, and Rafelski 2015; Huet et al. 2018). In order to create strains amenable to mitochondrial volume analysis, we transfected both ATPTG8Ty/cKD and ATPTG9Ty/cKD parasite strains with plasmids encoding GFP fused to the mitochondrial targeting signal of SOD2 and TdTomato fused to the inner membrane complex protein IMC1 (Harding et al. 2016; Pino et al. 2007). As *T. gondii* mitochondrial morphology changes dramatically during endodyogeny (Nishi et al. 2008), the IMC1-TdTomato signal allowed us to exclude any parasites undergoing division from our analysis. Using z-stacking and the MitoGraph program (Viana, Lim, and Rafelski 2015), we were able to quantify mitochondrial volume and observe 3D changes in mitochondrial ultrastructure from SOD2-GFP signal as ATPTG8Ty/cKD and ATPTG9Ty/cKD parasites were treated with ATc over 72 hours. With this approach, we also observed mitochondrial fragmentation in parasites treated with ATc from both conditional knockdown strains (Figure 3B). Additionally, quantification of mitochondrial volumes from 100 vacuoles per timepoint illustrated that after just 24 hours of ATc treatment, mitochondrial volume decreased significantly in both strains and continued to decline at 48 and 72 hour timepoints as well (Figure 3C). These results, in combination with our observations via immunofluorescence, confirm that mitochondrial fragmentation and volume loss occur upon ATPTG8 and ATPTG9 knockdown.

### Knockdown of ATP synthase-associated CHCH domain proteins causes decreased cristae density

To continue uncovering the mitochondrial ultrastructure phenotypes observed upon CHCH domain protein knockdown, we next wanted to investigate whether knockdown led to changes in mitochondrial cristae. Cristae are invaginations of the inner mitochondrial membrane associated with respiration efficiency and mitochondrial health (Quintana-Cabrera et al. 2018). In both yeast and mammals, the ATP synthase dimer has been shown to be crucial in shaping and maintaining mitochondrial cristae (Blum et al. 2019; Davies et al. 2012). A similar phenomenon has previously been observed in *T. gondii*, in which loss of ATP synthase integrity or its ability to assemble into higher order structures have both disrupted proper cristae formation (Huet et al. 2018; Muhleip et al. 2021). We utilized transmission electron microscopy to observe changes in cristae density between the parental strain treated with ATc and each conditional knockdown strain treated with ATc or vehicle control for 72 hours. While there were some variations between the parental and ATPTG8Ty/cKD or ATPTG9Ty/cKD strains -ATc, there was a significant decrease in cristae density when ATc was added to ATPTG8Ty/cKD or ATPTG9Ty/cKD parasites for 72 hours (Figure 4A,B). Similar mitochondrial areas were investigated for all samples, except for ATPTG9Ty/cKD (Figure S2). Overall, the depletion of ATPTG8 and ATPTG9 causes mitochondrial fragmentation, loss of mitochondrial volume, and decreased cristae density.

**Figure 4:**
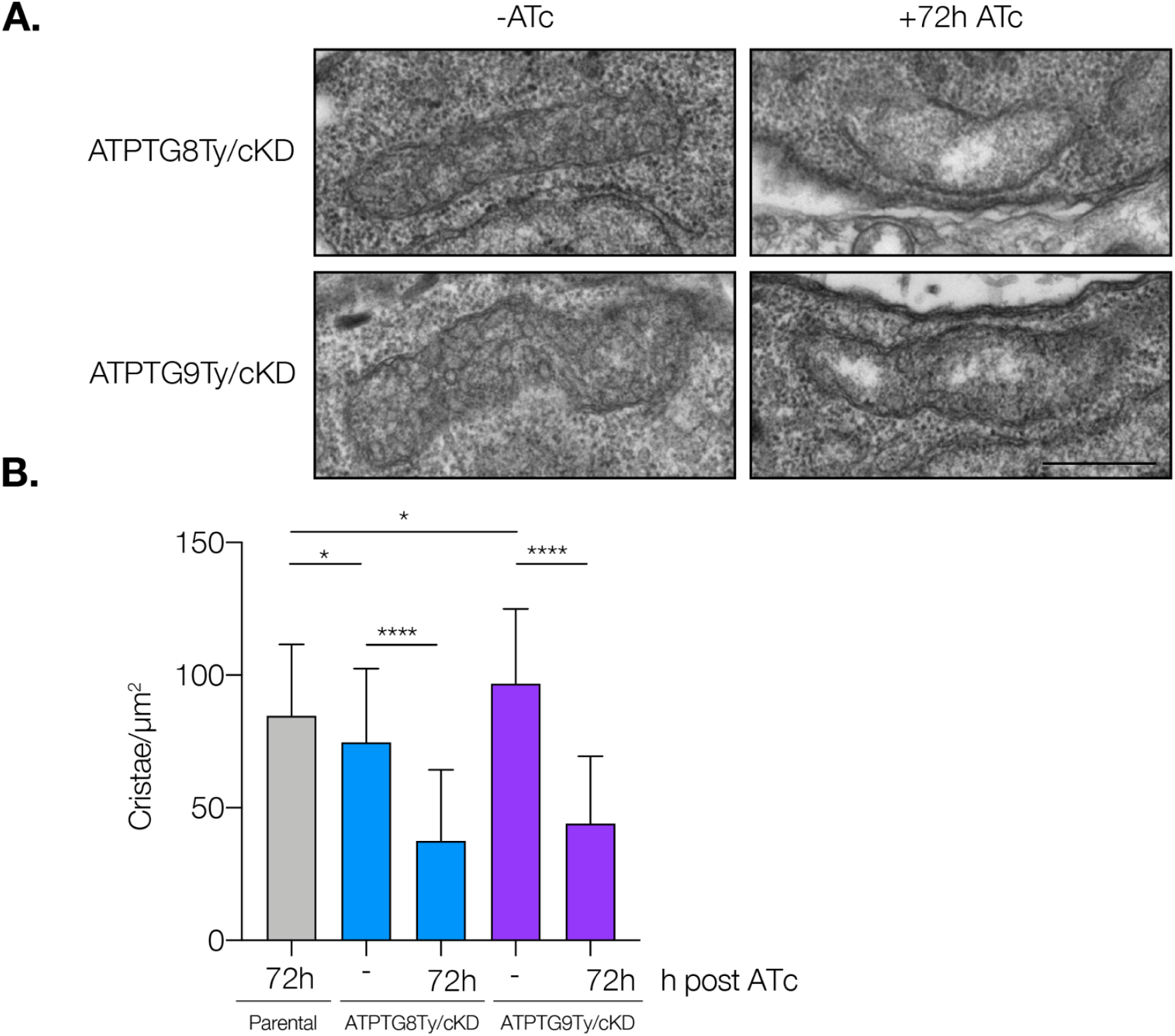
Mitochondrial cristae density decreases significantly upon CHCH-domain protein knockdown. Mitochondrial cristae density of parental, ATPTG8Ty/cKD, and ATPTG9Ty/cKD parasites treated with ATc or vehicle control (-) was determined using transmission electron microscopy. **A**. Representative electron micrographs of mitochondria from ATPTG8Ty/cKD and ATPTG9Ty/cKD parasites treated with ATc or vehicle control (-) for 72h. Scale bar: 500nm. **B**. Quantification of cristae/µm^2^ of mitochondrial area of parental parasites +72h ATc and ATPTG8Ty/cKD or ATPTG9Ty/cKD treated with ATc or vehicle control (-) for 72h. Data represent mean ± SD for 60 sections of each condition, which were blinded prior to analysis. Unpaired, two-tailed t-test (p < 0.0001: ****, p = 0.01 to 0.05: *).

### CX_9_C cysteine residues are required for mitochondrial localization of ATP synthase-associated CHCH domain proteins

A defining characteristic of CHCH domain proteins is their cysteine residues: two pairs of cysteine residues each separated by nine other amino acids in separate α-helices (Modjtahedi et al. 2016). Upon import into the mitochondrial intermembrane space (IMS), the CHCH domain cysteines are oxidized to form disulfide bonds. These intramolecular bonds have been shown to be critical for the proper folding, stability, and import of the proteins into the IMS (M. Darshi et al. 2012; Bourens et al. 2012; Aras et al. 2015). To investigate whether the cysteine residues in ATPTG8 and ATPTG9 were similarly important, we created plasmids containing either an HA-tagged wildtype coding sequence of each gene, or an HA-tagged copy in which cysteine residues of the CHCH domain were mutated to serines. These plasmids were then transfected into the respective ATPTG8Ty/cKD or ATPTG9Ty/cKD strain (Figure 5A). Within 24 hours of transfection, immunofluorescence assays were utilized to observe the localization of the HA-tagged wildtype and cysteine∆serine CHCH domain proteins (Figure 5B). While the exogenously expressed HA-tagged wildtype copy of each gene clearly localized to the parasite mitochondrion, the mutant proteins mis-localized and were instead found in the cytoplasm (Figure 5B). Thus, the cysteine residues in the CHCH domains of both ATPTG8 and ATPTG9 are critical for mitochondrial localization of both proteins.

**Figure 5:**
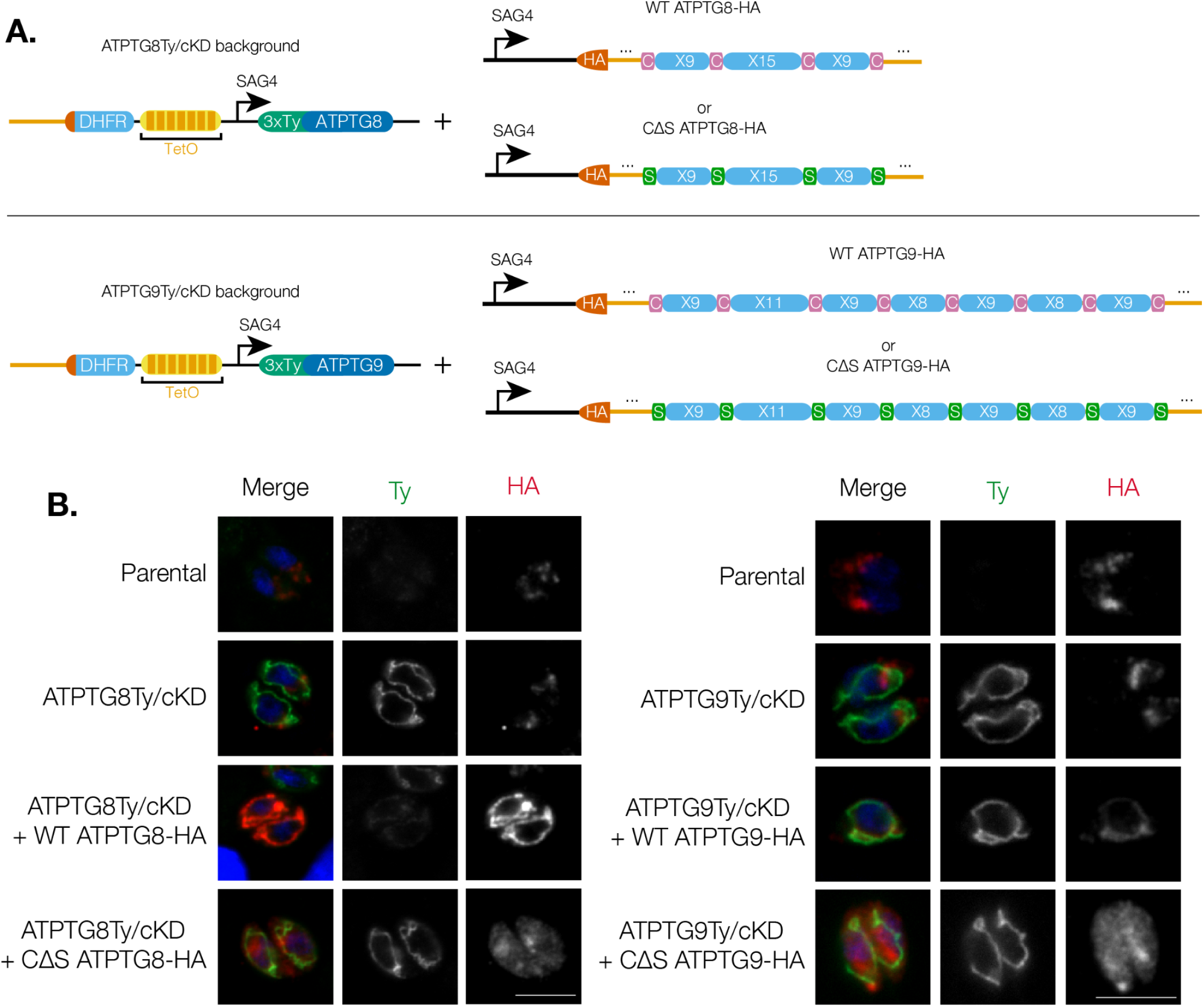
CX_9_C cysteine residues are necessary for mitochondrial localization of ATP synthase-associated CHCH domain proteins. **A**. Schematic of the strategy to transiently express exogenous HA-tagged wildtype (WT) or mutant CHCH cysteineΔserine (CΔS) copies of ATPTG8 or ATPTG9 in the respective tagged conditional knockdown strain. **B**. Immunofluorescence assays of parasites from TATi/Δku80 parental, ATPTG8Ty/cKD or ATPTG9Ty/cKD strains transiently expressing the HA-tagged WT or CΔS copy of the respective gene, as well as un-transfected control parasites. Intracellular parasites were fixed and stained for Ty (green) and HA (red). Scale bar: 5µm. Images are representative of 3 independent replicates.

## DISCUSSION

While the mitochondrion is commonly known as “the powerhouse of the cell,” the source of its power stems from the ATP synthase: the multisubunit enzymatic complex that generates large amounts of cellular energy in the form of ATP. The subunit composition of the ATP synthase is remarkably well-conserved among a wide range of phyla, including yeast and mammals (Jonckheere, Smeitink, and Rodenburg 2012). While the *T. gondii* ATP synthase contains most of the canonical subunits, it also contains 17 other largely uncharacterized, apicomplexan-specific subunits. In the present study, we characterized two novel subunits of the *T. gondii* ATP synthase that contain CHCH domains: ATPTG8 and ATPTG9.

Using conditional gene knockdown systems, we validated that both genes are essential for the *T. gondii* lytic cycle, as was predicted by a genome-wide CRISPR screen (Sidik et al. 2016). Furthermore, we were able to determine that both genes are critical for ATP synthase structural stability as well as function. Though oxidative phosphorylation significantly decreased, knockdown of either CHCH domain protein had no effect on the ability of the parasites to perform glycolysis, illustrating their metabolic flexibility. These observations of ATP synthase destabilization and decreased oxidative phosphorylation fit with findings in yeast, in which many CHCH domain proteins are found to play a role in the biogenesis and function of respiratory chain complexes (S. Longen et al. 2009; Cavallaro 2010). It has been hypothesized that most CHCH domain proteins act as structural scaffolds for large mitochondrial complexes. This may be due to the CHCH domain motif serving as a stable, yet easily modifiable, building block that can be folded through a conserved mechanism and regulated by mitochondrial REDOX state (Cavallaro 2010). While CHCH domain proteins have never previously been observed as subunits of the ATP synthase in any other organism, ATPTG8 and ATPTG9 likely provide additional structural support to the exceptionally large *T. gondii* ATP synthase complex.

Although *T. gondii* tachyzoites can continue to generate ATP via glycolysis when ATP synthase function is disrupted, we also demonstrated that ATP synthase structural stability is critical for maintaining proper mitochondrial membrane potential. Previous studies in mouse neurons illustrated that when OSCP, an ATP synthase subunit involved in stabilization of the complex, was downregulated, the cells experienced membrane potential depolarization (Beck et al. 2016). The destabilization of the complex eventually leads to the dissociation of the F1 portion and the formation of a channel within the c ring of the ATP synthase, which allows ions to leak through (Beck et al. 2016). It is possible that downregulation of ATPTG8 and ATPTG9 could destabilize the *T. gondii* ATP synthase in a way that allows ion leak across the inner mitochondrial membrane, thus resulting in the observed membrane potential depolarization.

Additionally, we found that knockdown of both genes caused mitochondrial fragmentation and a sharp decrease in mitochondrial volume. Similar mitochondrial defects in yeast have been observed when three different CHCH domain proteins were knocked out, leading to swollen, irregular, shorter, or less interconnected mitochondrial tubules (S. Longen et al. 2009). Furthermore, we showed that knockdown of both ATPTG8 and ATPTG9 resulted in significant reduction of mitochondrial cristae density. The bulbous mitochondrial cristae in *T. gondii* are shaped by pentagonal pyramid ATP synthase complexes (Muhleip et al. 2021), in contrast to the mammalian lamellar cristae that are shaped by a large ATP synthase dimer angle (Blum et al. 2019). Our work adds to a growing volume of evidence illustrating that disrupting both the ATP synthase dimer or pentagonal pyramids results in a loss of cristae integrity in *T. gondii* (Huet et al. 2018; Muhleip et al. 2021).

Though our study illustrates that CHCH domain proteins play important roles in the *T. gondii* mitochondrion, the mechanism by which these proteins are imported and stabilized within the *T. gondii* mitochondrion remains unclear. In yeast, a CHCH domain protein termed Mia40 (or CHCHD4 in humans) resides in the IMS and assists in the oxidative folding of other CHCH domain proteins following their mitochondrial import. Interactions between CHCH domain proteins and Mia40 are mediated by an intermembrane targeting sequence (ITS). The ITS consists of critical hydrophobic residues in an amphipathic helix either upstream or downstream of one of the CHCH domain cysteine residues, and the majority of CHCH domain proteins lack canonical mitochondrial matrix targeting signals (Cavallaro 2010; Sideris et al. 2009). Although ATPTG8 and ATPTG9 are not predicted to contain matrix targeting signals, they contain putative ITS motifs. Tagging either gene at the N terminus did not affect their mitochondrial localization, confirming their lack of matrix targeting signals. Ultimately, through its interaction with imported CHCH domain proteins, Mia40 is able to oxidize key cysteine residues, resulting in the formation of the two disulfide bonds (Peleh, Cordat, and Herrmann 2016). Oxidized Mia40 is then regenerated through interaction with the flavin-linked sulfhydryl oxidase Erv1 in yeast (or ALR in humans) (Ang et al. 2014; Mesecke et al. 2005).

Intriguingly, many protozoan parasites, including apicomplexans, express CHCH domain proteins but seem to lack Mia40 homologs (Mallo et al. 2018). It has been hypothesized that Erv1 may have functioned independently as part of an ancestral import pathway for CHCH domain proteins in early eukaryotic lineages (Allen, Ferguson, and Ginger 2008). However, complementation studies attempting to replace yeast Mia40 with protozoan Erv1 have failed to validate this (Specht et al. 2018). While three putative Erv1 homologs have been identified in the *T. gondii* genome (van Dooren et al. 2016), the identity of the oxidoreductase responsible for folding CHCH domain proteins in these parasites remains unknown.

Our work demonstrates that the cysteine residues within ATPTG8 and ATPTG9 are critical for their proper mitochondrial localization; while transient expression of wildtype ATPTG8 or ATPTG9 showed consistent mitochondrial localization, cysteine∆serine copies of both genes appeared as diffuse cytoplasmic signal. These observations are supported by previous studies in which CHCH domain cysteine mutants appear to exhibit disrupted localization, co-localize with lysosomes, or be degraded via other methods (Manjula Darshi et al. 2011; Bourens et al. 2012; Aras et al. 2015). These data emphasize the importance of the CHCH domain cysteines in the localization and stability of ATPTG8 and ATPTG9 and further suggest that *T. gondii* CHCH domain proteins must interact with an yet to be determined oxidoreductase to be properly folded and maintained within the mitochondrion. As the putative parasite oxidoreductase likely diverges significantly from its human counterpart, it could represent another novel therapeutic target and its identification warrants future study.

Additionally, while there appears to be a great deal of conservation in terms of ATP synthase subunit composition between *T. gondii* and the related apicomplexans of the genus *Plasmodium spp*., very little is known about the ATP synthase in this group of malaria-causing parasites. As the ATP synthase has been shown to be essential for survival of the insect stages of *Plasmodium berghei* (Sturm et al. 2015), drugs targeting this complex could block malaria transmission without affecting the mammalian host. However, additional characterization of this important complex in *Plasmodium spp*. is needed.

In summary, our work provides the first insight into the roles played by ATP synthase-associated CHCH domain proteins and increases the depth of knowledge surrounding the divergence of the apicomplexan ATP synthase. Future work remains to characterize other putative CHCH domain proteins in *T. gondii* (Table 1), determine the identity of the oxidoreductase involved in their mitochondrial import, and elucidate potential regulatory mechanisms of this important class of proteins. Because novel drugs against these parasites are needed to combat constantly evolving resistance against current therapeutics, characterizing essential, divergent parasite proteins can offer new opportunities for future drug discovery.

**Table 1:** Putative CHCH domain proteins in *T. gondii*. List of CHCH domain proteins primarily generated via ToxoDB BLASTp analysis based on datasets of CHCH domain proteins in other organisms (Cavallaro 2010; Modjtahedi et al. 2016). LOPIT data: Barylyuk et al. 2020. CRISPR scores: Sidik et al. 2016.

| Gene ID | ToxoDB annotation | CRISPR Score | LOPIT | Protein size (aa) | CHCH domain | Notes |
| --- | --- | --- | --- | --- | --- | --- |
| TGGT1_203460 | hypothetical protein | -3.5 | Mitochondrion-soluble | 119 | C9C17C9C (51-89) | Homology to TRIAP1/Mdm35, a protein involved in mitochondrial lipid homeostasis (BLASTp E: 2e-05) |
| TGGT1_208300 | hypothetical protein | -1.9 | Mitochondrion-soluble | 105 | C9C10C9C (65-96) |  |
| TGGT1_213940 | CHCH domain-containing protein | -2.2 | Mitochondrion-soluble | 149 | C9C9C9C (111-141) | Homology to human CHCHD2/CHCHD10 (BLASTp E: 0.002) |
| TGGT1_240550 | putative copper chaperone COX17-1 | -2.7 | Mitochondrion-soluble | 77 | C9C8C9C (39-68) |  |
| TGGT1_254260 | COX19 cytochrome c oxidase assembly family protein | -3.3 | Mitochondrion-soluble | 256 | C9C32C9C (29-82) |  |
| TGGT1_258060 | putative myosin heavy chain | -4.1 | Mitochondrion-membranes | 208 | C9C15C9C (129-165) | Identified as subunit of ATP synthase by this study, Huet et al., 2018, Muhleip et al., 2021, Maclean et al., 2021 |
| TGGT1_268740 | hypothetical protein | -1.6 | Mitochondrion-soluble | 187 | C9C10C9C (91-122) |  |
| TGGT1_285510 | hypothetical protein | -1.9 | Mitochondrion-membranes | 166 | C9C11C9C8C9C8C9C (14-84) | Identified as subunit of ATP synthase by this study, Huet et al., 2018, Salunke et al., 2018, Muhleip et al., 2021, Maclean et al., 2021 |
| TGGT1_286070 | hypothetical protein | -3.4 | Mitochondrion-soluble | 86 | C9C29C9C (18-68) |  |
| TGGT1_290710 | hypothetical protein | 0.7 | Not localized | 236 | C9C9C11C** (193-225) | Identified as subunit of ATP synthase by Muhleip et al., 2021 and Maclean et al., 2021. **Does not contain a typical twin Cx9C motif |
| TGGT1_297465 | hypothetical protein | 0.7 | Not localized | 94 | C9C19C9C (27-67) |  |
| TGGT1_311390 | tRNA (guanine(9)-N(1))-methyltransferase | 1.4 | Not localized | 605 | C9C9C9C (317-347) |  |
| TGGT1_313160 | hypothetical protein | -4 | Not localized | 138 | C9C11C9C (51-83) |  |
| TGGT1_320140 | putative ubiquinol-cytochrome c reductase hinge protein | -3.7 | Mitochondrion-membranes | 89 | C9C13C9C (41-75) | Identified as CIII subunit QCR6/Hinge protein in Hayward and van Dooren, 2019 and Maclean et al., 2021 |

## ACKNOWLEDGMENTS

We would like to acknowledge Wandy Beatty at the Washington University Molecular Microbiology Imaging Facility for acquiring transmission electron microscopy images. Additionally, we would like to thank Julie Nelson from the UGA Cytometry Shared Resource Laboratory as well as Dr. Muthugapatti Kandasamy from the UGA Biomedical Microscopy Core for their expertise and use of their equipment. Feedback, antibodies, and plasmids courtesy of Dr. Silvia Moreno’s laboratory were integral in completing this work. Further, we would like to thank Dr. Alex Rosenberg for feedback on this manuscript. Lastly, we would like to thank Brittany Henry and Kaelynn Parker for experimental assistance.

## MATERIALS AND METHODS

### Parasite culture

RH/TATi/Δku80 (Sheiner et al. 2011) tachyzoites and their derivatives were maintained in human foreskin fibroblasts (HFFs) (ATCC, cat. no. SCRC-1041). Strains were cultured at 37ºC and 5% CO_2_ in DMEM supplemented with 2mM glutamine (GeminiBio, cat. no. 400-106) and 3% heat-inactivated fetal calf serum (IFS).

### Plasmid generation

To generate the ATPTG8Ty/cKD and ATPTG9Ty/cKD strains, the pU6-Universal plasmid (Addgene, cat. no. 52694) was digested using the BsaI restriction enzyme then sgRNAs targeting the N terminus of each gene (P1 and P2 for ATPTG8 or P5 and P6 for ATPTG9) were inserted into the plasmid using Gibson assembly. A repair template encoding a pyrimethamine-resistant copy of the dihydrofolate reductase (DHFR) cassette, the *T. gondii* SAG4 promoter, a tetracycline-inducible operator, and a 3xTy epitope tag was amplified from the DHFR-SAG4-TetO7-3xTy plasmid (a kind gift from Silvia Moreno) using P3 and P4 for ATPTG8Ty/cKD or P7 and P8 for ATPTG9Ty/cKD.

To modify the ATPTG8Ty/cKD strain so that it would transiently express either wildtype or cysteine∆serine copies of ATPTG8, a plasmid that contained the SAG4 promoter, an N-terminal HA tag, the ATPTG8 CDS, and 1000bp of the endogenous ATPTG8 3’ UTR (SAG4-HA-ATPTG8WT) was assembled via Gibson assembly. To create a plasmid encoding a cysteine∆serine mutant copy of the gene (SAG4-HA-ATPTG8CtoS), a portion of the ATPTG8 CDS was digested out with EcoNI and StuI. An oligonucleotide carrying the portion of the ATPTG8 CDS with all cysteine to serine mutations (P9) was amplified with P10 and P11 and inserted using Gibson assembly.

A similar approach was used for ATPTG9: a plasmid containing the SAG4 promoter, an N-terminal HA tag, the ATPTG9 CDS, and 1000bp of the endogenous ATPTG9 3’ UTR (SAG4-HA-ATPTG9WT) was assembled via Gibson assembly. To create a plasmid encoding a cysteine∆serine mutant copy of the gene (SAG4-HA-ATPTG9CtoS), a portion of the ATPTG9 CDS was digested out with KpnI and EcoRI. An oligonucleotide carrying the portion of the ATPTG9 CDS with all cysteine to serine mutations (P14) was amplified with P15 and P16 and inserted using Gibson assembly.

### Parasite strain generation

To create the ATPTG8Ty/cKD and ATPTG9Ty/cKD strains, TATi/Δku80 parasites were transfected as previously described (Sidik et al. 2014). These transfections were conducted with 25-50µg of pU6-Universal plasmid encoding Cas9 and an sgRNA targeting the N terminus of each gene. The repair oligonucleotide co-transfected with this plasmid encoded a pyrimethamine-resistant DHFR cassette, the *T. gondii* SAG4 promoter, tetracycline-inducible operator, and an in-frame 3xTy epitope tag flanked on either end by 40bp of homology to the 5’ UTR or N terminus of the gene. Pyrimethamine (Sigma Aldrich, cat. no. 46706-250MG) was used at 3µM to select for parasites containing the integration. Expression of the Ty tag and its downregulation after addition of 0.5µg/ml anhydrotetracycline (ATc) were confirmed via immunofluorescence assay and western blot. Positive integrants were subcloned via serial dilution to obtain a monoclonal population.

For mitochondrial volume analysis experiments, both the ATPTG8Ty/cKD and ATPTG9Ty/cKD strains were transfected with pT8mycSOD2(SPTP)GFPmycHX (Pino et al. 2007) and TubIMC1TdTomato-CAT plasmids (Harding et al. 2016). Double positive parasites were sorted via FACS to isolate clonal populations stably expressing both fluorescent proteins.

For localization of exogenously expressed wildtype or cysteine∆serine ATPTG8 or ATPTG9, ∼50-100µg of SAG4-HA-ATPTG8WT or SAG4-HA-ATPTG8CtoS plasmids were transfected into the ATPTG8Ty/cKD line, while ∼50-100µg of SAG4-HA-ATPTG9WT or SAG4-HA-ATPTG9CtoS plasmids were transfected into the ATPTG9Ty/cKD line. Immediately after transfection, 40µl of parasites were added to coverslips pre-seeded with HFF cells for monitoring of transient HA expression within 24 hours of transfection.

### Plaque assays

To determine effects of CHCH domain protein knockdown on the *T. gondii* lytic cycle, 500 TATi/Δku80 parental, ATPTG8Ty/cKD, or ATPTG9Ty/cKD parasites were added to each well of a 6-well plate pre-seeded with HFF cells and left undisturbed for 7-8 days. Wells were treated with 0.5µg/ml ATc or ethanol (vehicle control) for the entirety of the experiment. For the recovery assays, parasites were pre-treated with ATc or vehicle control for 72 hours prior to being added to the wells without ATc or vehicle control. After 7-8 days, wells were washed with PBS, fixed in 95% ethanol for 10 minutes then stained with a crystal violet solution (2% crystal violet, 0.8% ammonium oxalate, 20% ethanol) for 5 minutes. Wells were subsequently washed again then scanned for analysis.

### Western blotting

To prepare samples for western blotting, pellets containing approximately 2e7 parasites were resuspended in 2X Laemmli buffer (20% glycerol, 5% 2-mercaptoethanol, 4% SDS, 0.02% bromophenol blue, 120 mM Tris-HCl pH 6.8) then boiled at 100°C for 5 minutes. Following separation by SDS-PAGE, proteins were transferred to nitrocellulose membranes and probed with mouse anti-Ty, mouse anti-tubulin (Developmental Studies Hybridoma Bank at the University of Iowa, cat. no. 12G10), or rabbit anti-F1β (Agrisera, cat. no. AS05 085). For signal detection using an Odyssey infrared imager (LI-COR Biosciences), donkey anti-mouse IgG conjugated to IRDye 800CW (VWR, cat. no. 102673-332) or goat anti-rabbit IgG conjugated to IRDye 680RD (VWR, cat. no. 102673-410) were used as secondary antibodies.

### Blue native polyacrylamide gel electrophoresis (BN-PAGE)

For BN-PAGE, 2×10^7^ parasites were solubilized with 2.5% digitonin in a solution containing 1X NativePAGE sample buffer (Thermo Fisher Scientific, cat. no. BN2008). To accurately estimate the molecular weight of large membrane-bound complexes, 50µg of bovine heart mitochondria (Abcam, cat. no. ab110338) were solubilized under the same conditions as the parasite samples for use as a standard (Evers et al. 2021). Following separation on a NativePAGE 3-12% Bis Tris protein gel (Thermo Fisher Scientific, cat. no. BN1001BOX), proteins were transferred to a PVDF membrane and incubated with rabbit anti-F1β. Membranes were then incubated with goat anti-rabbit IgG conjugated to HRP (VWR, cat. no. 102645-182) secondary antibody. After incubation with enhanced chemiluminescence (ECL) substrate (VWR, cat. no. PI32209), autoradiography film (MTC Bio, cat. no. A8815) was exposed to the membrane and developed on an X-ray film processor.

### Cellular ATP concentration measurements

To measure the ATP concentration of the parasites, HFFs were infected with either parental, ATPTG8Ty/cKD, or ATPTG9Ty/cKD strains and treated with vehicle control (ethanol) or 0.5µg/ml ATc for 72h. As previously described (Huet et al. 2018), the monolayer was first washed with PBS to remove any extracellular parasites. Fluorobrite DMEM (Thermo Fisher Scientific, cat. no. A1896701) containing 1% IFS and HALT protease inhibitors (VWR, cat. no. PI78440) was added to the cells containing intracellular parasites before they were scraped and syringe-released. Following removal of host cell debris, parasites were pelleted, washed in carbon-free DMEM (Fisher Scientific, cat. no. A1443001), then resuspended in carbon-free DMEM to a final concentration of 6×10^6^ parasites/ml. To determine the initial ATP concentration for each strain, 50µl of each parasite solution was added to the wells of a 96-well PCR plate with 50µl carbon-free DMEM before immediately being flash frozen in liquid nitrogen. Additionally, 50µl of each parasite solution was added to a plate along with equal amounts of the following compounds at the listed final concentrations: 5mM 2-deoxyglucose (Sigma Aldrich, cat. no. D6134-5G) + 25mM glucose (Sigma Aldrich, cat. no. G7021-100G) or 5mM 2-deoxyglucose + 2mM glutamine (Sigma Aldrich, cat. no. G8540-100G). Parasites were allowed to incubate with each solution for 1 hour at 37°C, 5% CO_2_ before being flash frozen. To determine the ATP concentration in each condition, 100µl of CellTiter-Glo reagent (Promega, cat. no. G7572) was added to the wells while samples thawed at room temperature for 1 hour. After thawing, 100µl of each sample was added to a white, flat-bottom 96-well plate then measured using a Molecular Devices SpectraMax i3x microplate reader. All samples and conditions were conducted in triplicate and ATP levels for each strain were normalized to the initial concentration. Six biological replicates were obtained from the parental strain and 3 replicates were obtained for ATPTG8Ty/cKD and ATPTG9Ty/cKD strains.

### Immunofluorescence assays

For the immunofluorescence assays, glass coverslips pre-seeded with HFFs were infected with approximately 40µL of freshly lysed parasites. While the parasites were intracellular, cells were fixed in 4% paraformaldehyde for 15 min at 4°C. Following fixation, cells were permeabilized for 8 minutes using a solution of 0.25% Triton X-100, then blocked for 10 minutes in a solution of PBS with 5% heat-inactivated fetal bovine serum (IFS) and 5% normal goat serum (NGS). Cells were then stained with mouse anti-Ty (Bastin et al. 1996), rabbit anti-Tom40 (a kind gift from Giel van Dooren), or rabbit anti-HA (Abcam, cat. no. ab9110) primary antibodies for 1 hour. Subsequently, cells were stained with Alexa-488-conjugated goat-anti-mouse (Invitrogen, cat. no. A32723) and Alexa-647-conjugated goat-anti-rabbit (Invitrogen, cat. no. A32733) secondary antibodies. Hoechst (Santa Cruz Biotechnology, cat. no. sc-394039) was used to stain cell nuclei. Coverslips were mounted onto slides with Prolong Diamond (Thermo Fisher, cat. no. P36961). Images were acquired using an ECHO Revolve microscope and the ECHO Pro application. Image analysis and processing were conducted using Fiji, Adobe Photoshop 2022, and Adobe Illustrator 2022.

### Flow Cytometry

To quantify membrane potential changes when ATPTG8 or ATPTG9 genes were knocked down, intracellular tachyzoites from parental, ATPTG8Ty/cKD or ATPTG9Ty/cKD strains were treated with ATc or vehicle control (ethanol) for 72 hours then stained with 50nM MitoTracker DeepRed (Life Technologies, cat. no. M22426) for 1 hour at 37ºC and 5% CO_2_. For FCCP and vehicle control (DMSO) samples, intracellular parental tachyzoites were stained with a solution containing 50nM MitoTracker DeepRed (Life Technologies, cat. no. M22426) and 10µM carbonyl cyanide 4-(trifluoromethoxy)phenylhydrazone (FCCP) (Sigma Aldrich, cat. no. C2920-10MG) or an equivalent amount of DMSO (vehicle control). After 1 hour, parasites were syringe-released, filtered, and pelleted before washing once with PBS (10µM FCCP or DMSO were included in PBS for the FCCP and DMSO samples). Samples were resuspended in PBS or PBS containing 10µM FCCP or DMSO for analysis via flow cytometry. Membrane potential was analyzed on an Agilent NovoCyte Quanteon using a 637nm laser and emission was measured using a 695/40 filter.

### Mitochondrial volume analysis

HFFs seeded in 35mm glass-bottom microscopy dishes were infected with ATPTG8Ty/cKD or ATPTG9Ty/cKD parasites expressing SOD2-GFP and IMC1-TdTomato. Parasites were treated with ethanol (vehicle control) for 72h, or 0.5µg/ml ATc for 24, 48, or 72h. Images of vacuoles containing either one or two parasites were acquired using the 60x lens of a DeltaVision II Microscope System II. Each vacuole was imaged using a Z-stack of 25 images spaced 0.2µm apart such that the top and bottom images of the stack showed slightly out-of-focus mitochondria. Vacuoles containing parasites undergoing endodyogeny, as observed by the appearance of daughter cells using the IMC1-TdTomato signal, were excluded from analysis. Images were analyzed and mitochondrial volume was calculated using the MitoGraph system as previously described (Viana, Lim, and Rafelski 2015). At least 100 vacuoles were analyzed over 2 or 3 biological replicates for each strain.

### Transmission Electron Microscopy

In order to quantify changes in mitochondrial cristae density upon CHCH domain subunit knockdown, HFF cells infected with TATi/Δku80 +ATc 72h, ATPTG8Ty/cKD -ATc 72h, ATPTG8Ty/cKD +ATc 72h, ATPTG9Ty/cKD -ATc 72h, or ATPTG9Ty/cKD +ATc 72h parasites were trypsinized, pelleted, then fixed in a 50mM phosphate (Sigma Aldrich cat. no. P5655-100G; Sigma Aldrich cat. no. P9666-100G) buffer containing 1% glutaraldehyde (Fisher Scientific. cat. no. NC1536477) and 1% OsO_4_ (Fisher Scientific, cat. no. NC9402523) for 45 minutes at 4ºC. Samples were washed three times in cold 50mM phosphate buffer, then rinsed in cold dH_2_O. Staining and imaging were conducted as previously described (Huet et al. 2018). En bloc staining was performed with 1% aqueous uranyl acetate (Ted Pella Inc.) at 4ºC for 3 hours. After rinsing with dH_2_O, a series of ethanol solutions was used to dehydrate the samples before they were embedded in Eponate 12 resin (Ted Pella Inc.). 95 nm sections were obtained using a Leica Ultracut UCT ultramicrotome (Leica Microsystems Inc.) before staining with lead citrate and uranyl acetate. Samples were viewed using a JEOL 1200 EX transmission electron microscope (JEOL USA Inc.) and images were captured using an AMT 8-megapixel digital camera and AMT Image Capture Engine V602 software (Advanced Microscopy Techniques). For cristae quantification, 60 sections of each strain that were pre-selected to contain parasite mitochondria were blinded. Fiji software was utilized to measure mitochondrial area and cristae were counted manually. Images were subsequently un-blinded and a student’s t-test was utilized to determine differences in cristae density and mitochondrial area between strains.

**Supplemental Figure 1:**
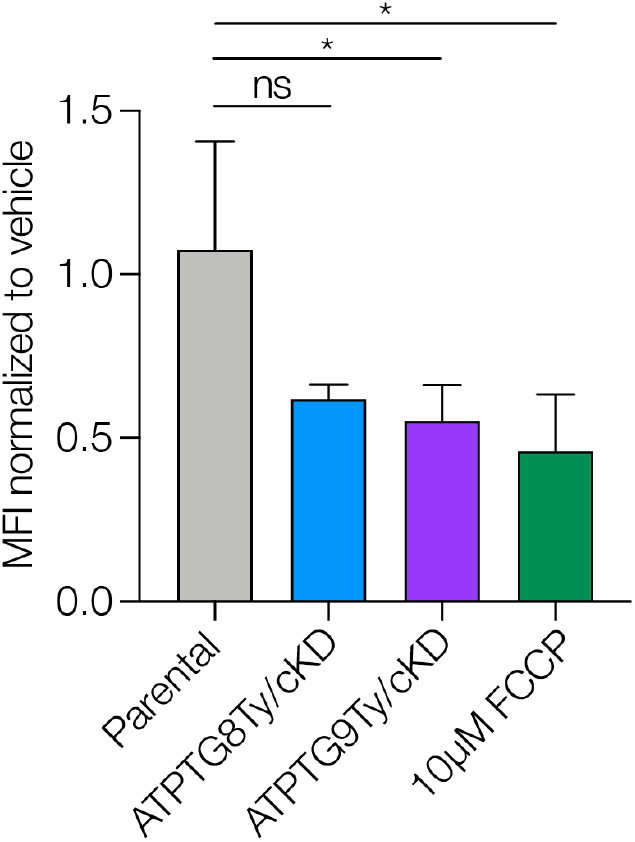
CHCH domain protein knockdown results in reduced mitochondrial membrane potential. Membrane potential of parental, ATPTG8Ty/cKD, and ATPTG9Ty/cKD parasites treated with ATc or vehicle control for 72h, or parental parasites treated with 10µM FCCP or vehicle control (DMSO) for 1 hour. Parasites were stained with 50nM MitoTracker and membrane potential was measured via flow cytometry. Each treatment condition was normalized to its vehicle control. Results represent mean ± SD for 3 independent replicates of ATPTG8Ty/cKD, ATPTG9Ty/cKD, and FCCP treatments or 4 independent replicates of parental strains. Unpaired, two-tailed t-test (ns = not significant, p = 0.01 to 0.05: *).

**Supplemental Figure 2:**
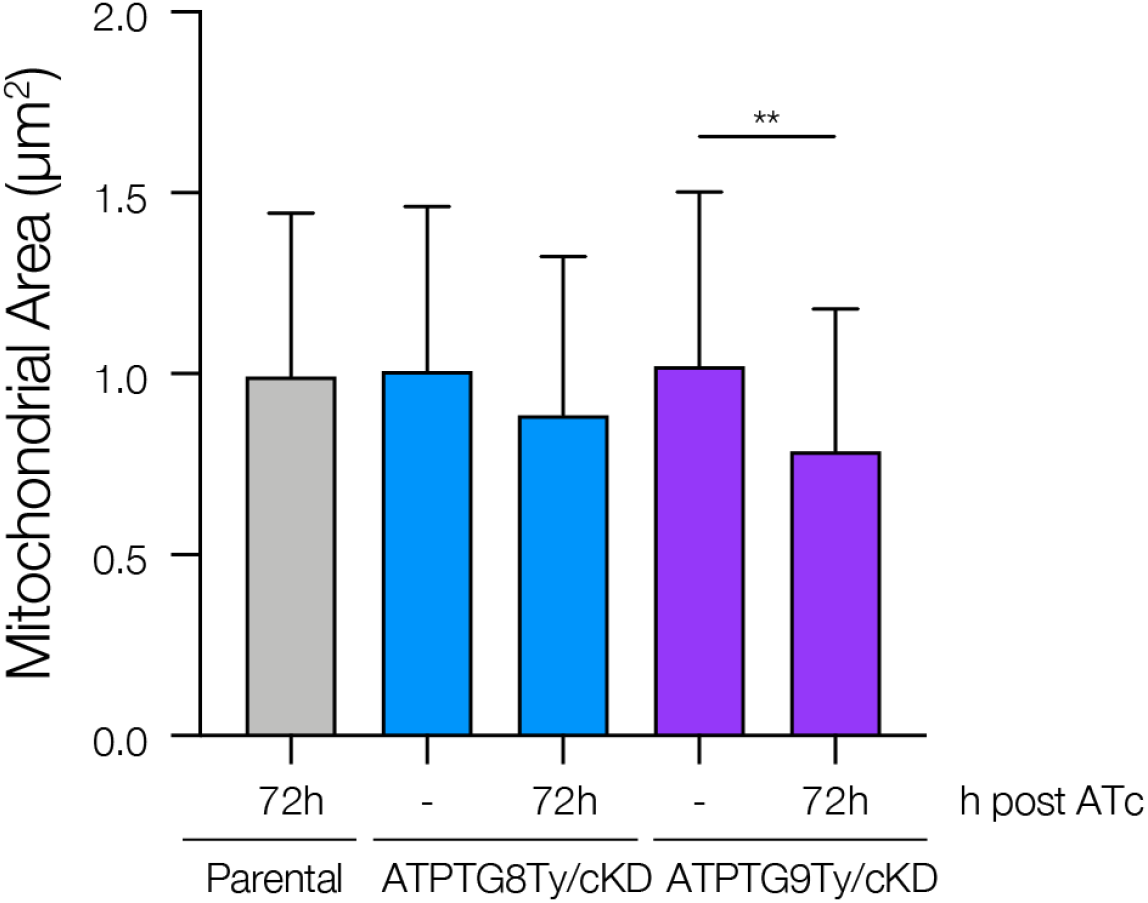
Mitochondrial areas measured during transmission electron microscopy analysis. Quantification of mitochondrial area (µm^2^) from parental parasites +72h ATc and ATPTG8Ty/cKD or ATPTG9Ty/cKD treated with ATc or vehicle control (-) for 72h as part of Figure 4. Data represent mean ± SD for 60 sections of each condition, which were blinded prior to analysis in Fiji. Unpaired, two-tailed t-test (p = 0.001 to 0.01: **).

## SUPPLEMENTARY MATERIAL

**Table S1: Summary of oligonucleotides used in the present study**.

